# Modeling asymmetric Thermal Performance Curves for proportion data

**DOI:** 10.1101/2025.10.17.683179

**Authors:** Leah R. Johnson, Catherine Lippi, Paul J. Huxley, Sadie J. Ryan

**Affiliations:** Virginia Tech, Statistics, Blacksburg, VA; Imperial College, Infectious Disease Epidemiology, London, UK; University of Florida, Geography, Gainesville, FL

## Abstract

Ectotherms are subject to temperature impacts on a variety of functional traits, such as fecundity and development. Quantifying the shape of the response of traits to temperature involves fitting (usually) non-linear curves to empirically derived data. This study introduces the LINEX Thermal Performance Curve (TPC), a novel non-linear predictor for modeling asymmetric thermal responses in biological traits. Unlike traditional symmetric TPCs, the LINEX function captures a wide range of asymmetric shapes (e.g., left- or right-skewed), making it particularly suitable for traits like survival or fecundity that often exhibit non-normal, bounded distributions. The study implements the LINEX TPC within a Bayesian framework using the `bayesTPC` R package. We demonstrate the model’s flexibility and utility through simulations and experimental data from arthropod thermal traits. Results highlight the LINEX TPC’s superior fit for asymmetric traits compared to standard quadratic models, as evidenced by improved performance metrics. The approach addresses challenges in estimating uncertainty in bounded data, providing a robust tool for ecological and evolutionary studies that require precise thermal sensitivity modeling.

## 1 Introduction

Ambient temperature impacts a variety of functional traits in ectotherms, such as fecundity, development, or survival [1]. Quantifying the impact of temperature on such traits is usually achieved through empirical, laboratory-based experimental studies. One approach is to hold organisms at an array of fixed temperatures and measure how focal traits respond at each temperature. Patterns of how such traits respond to these experimental temperature gradients are often quantified using Thermal Performance Curves [TPCs, 2,3,4]. These TPCs are usually functions that are non-negative (as measured traits are typically non-negative). They are often hump- or U-shaped, and may be either symmetric or asymmetric (see [4] and [5] for a range of examples).

Often the observed trait measurements (or the average of these) may reasonably be assumed to be roughly normally distributed at a particular temperature, with the mean at that temperature described by a TPC. For example, across moderate temperatures mosquitoes often lay many eggs at once, so these counts (or the average across times/individuals) may be approximately normal. In these cases, standard fitting approaches including non-linear least squares (NLLS) may be used to fit the TPCs to data [6]. Further, as long as the data may reasonably be assumed to be non-negative and likely unbounded from above, the general approach may still work. For example, if data are strictly positive, such as wing lengths, then the standard TPC can be used, but linked to other distributions, such as a truncated normal or gamma, in order to better describe the variability of the data around the assumed TPC.

However, this approach is not ideal for data that are also bounded within an interval. Specifically, proportion data, where the number of “successes” within a given number of trials are recorded, are poorly captured by the standard TPC framework. This is because two conditions must hold in order for these types of data to be approximated by a normal distribution. First the probability of success must not be “too close” to either 1 or zero. This is problematic under the normal/truncated normal TPC framework because we often specifically want the TPC to be close to these limits at particular temperatures. Further, the number of trials must be sufficiently large, with the sample size needing to be even higher the closer that the success probability is to either zero or 1. If these conditions do not hold, the point estimate of the mean may be reasonable, but any uncertainty bounds will be problematic. For example, if the sample size is small, the error may be poorly estimated by the normal distribution. Additionally, when the uncertainty is problematic in the TPC where probabilities are near 1 or 0, the estimated ranges may not correspond to appropriate probabilities (i.e., they include values greater than 1 or less than 0). Thus, while NLLS and other similar fitting approaches can be used to obtain point estimates of TPC parameters, and estimate the resulting mean response, the uncertainty bounds for any parameters and the associated TPC are not meaningful. In addition to the complications of fitting needs, TPCs, in practice, are often constrained by the number of measurements taken in an experiment, along both axes, meaning we wish to minimize the number of parameters [7].

An alternative approach is to fit binomial success data directly using generalized linear models (GLMs). In this approach, the data are recorded as the number of successes with the corresponding number of trials instead of as a proportion. A “link” between the modeled probability, formulated in terms of a linear model, and these binomial outcomes is assumed. This approach is flexible and straightforward; however, because of the requirement that the underlying model be linear in the parameters (e.g., lines, polynomial terms or similar), a standard GLM framework cannot capture the asymmetric patterns common in thermal ecology.

In this short note we introduce a simple extension to the standard GLM formulation of TPCs for binomial data using a straightforward non-linear predictor function. This predictor function is highly flexible, allowing for a range of shapes (concave up or down, left or right skewed), has only 3 parameters to estimate, and is appropriate for use with a binomial modeling approach. First, we introduce the function and describe the parameters and interpretation. Second, we show how to implement it as a TPC to fit trait data in a Bayesian framework using the package `bayesTPC` in R. This allows us to accommodate the non-linear function together with the appropriate link and likelihood. Third, we give examples of fitting the TPC to simulated data, to explore some subtleties inherent in analysis of these kinds of data. Fourth, we use this to fit real data from examples of arthropod thermal traits. Finally, we discuss the use of the function more generally.

## 2 Methods

### 2.1 The Generalized Linear Modeling framework

A Generalized Linear Model (GLM) is a modeling framework that builds on linear regression modelling, a foundational model of statistics. We can write a multiple linear regression (MLR) model mathematically as

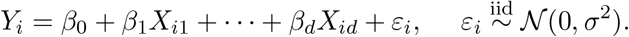

where *Y*_*i*_ is the *i*^th^ response variable, *X*_*i*1_ … *X*_*id*_ are the corresponding *d* predictor variables, the *β*s are the regression coefficients, and *ε* is the normally distributed noise with variance *σ*^2^. We can think of an MLR model as having three parts:

1. The ***systematic*** component: the explanatory variables are used to construct a linear predictor *η*_*i*_ = *β*_0_ + *β*_1_*X*_*i*1_ + · · · + *β*_*d*_*X*_*id*_.
2. The ***random*** component: *Y*_1_, …, *Y*_*n*_ are normally distributed with *Y*_*i*_ having mean *µ*_*i*_ and variance *σ*^2^.
3. The ***link*** between the random and systematic components: *µ*_*i*_ = *η*_*i*_, *i* = 1, …, *n*.

GLMs extend linear models by changing two requirements of the linear model. First, they allow different classes of distributions for the response variables. The most commonly used non-Normal distributions for GLMs are the Binomial (used for binary data) and the Poisson (used for positive counts). Second, they allow a more general link function:

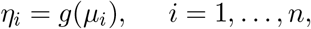

where *g* is a strictly increasing, twice differentiable function. Each family has a standard, or “canonical” link function. For example, the Poisson uses a log-link.

In this paper, we are interested in binary response data, that is data where the response *Y* is a binary variable (taking the value 0 or 1). In data on arthropod thermal traits, a common example is whether an egg that is laid successfully becomes an adult. Then a “success” (emergence of the adult) corresponds to the response *Y* = 1, and is *Y* = 0 to a failure. In this case we would use a logistic GLM such that the number of observed successes, *n*_*i*_, in a fixed number, *N*, trials is modeled as a Binomial random variable, and assuming a logit link to the linear predictor, that is:

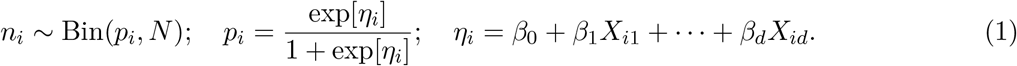

where again *η*_*i*_ is a linear predictor constructed as above.

In the standard GLM framework the linear predictor has two properties that are important. First, it has to give possible responses across the whole real line – that is when you plug in arbitrary values of predictors it can give values from −∞ to ∞. It is this property that necessitates the link function, which transforms the linear predictor to be within the necessary bounds of the probability model (between 0 and 1 for the binomial). The second property is that the predictor is linear in the *coefficients* of the predictor variables. Thus one can include polynomial terms, for example, because although one is taking powers of the variables, there are no powers (or other functions) of the coefficients in the predictor equation.

### 2.2 The LINEX TPC

From a theoretical perspective, it is possible to instead use a non-linear predictor – that is, one that fulfills the first requirement but not the second. Doing so means that other fitting methods must be used to fit the model. Because of this, we usually keep to the linear predictor unless it cannot produce a specific desired shape. In our case, we wish to be able to construct a predictor that is hump/u-shaped (like a quadratic) and asymmetric (unlike a quadratic) using as few parameters as possible.

One possible solution is to adapt an existing functional form. Linear Exponential (LINEX) loss functions are fairly common as constraints in the economic literature (e.g., [8]) and within the statistics literature as an alternative to quadratic loss when estimating model parameters (e.g., [9], [10], [11]). The functional form can be modified and used as a TPC in place of a linear predictor to generalize the GLM framework introduced above. We define the non-linear LINEX TPC *predictor* to be

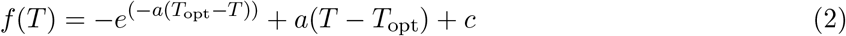

where *T*_opt_ determines the location of the optimum of the function, *c* − 1 is the height of the function at the maximum, and *a* governs a combination of the curvature (based on the magnitude of *a*) and skew. The direction of skew is determined by the sign of *a*: if *a >* 0 it is left skewed and if *a <* 0 it is right skewed (Fig 1, top and middle). The function can be modified slightly to enable a concave up shape, with the value of *a* still determining the skew,

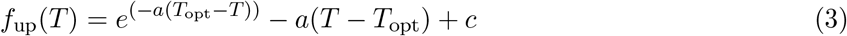

where the parameters are as for the concave down version, except that value at *T*_opt_ is now the *minimum* with a value *f*_up_(*T*_opt_) = *c* + 1 (Fig 1, bottom row). Either Eq 2 or Eq 3 can take the place of the linear predictor in the logistic model (Eq 1). We show the inverse logit transform of these equations (corresponding to the probability of success) in the right-hand panels of Fig 1.

**Figure 1.**
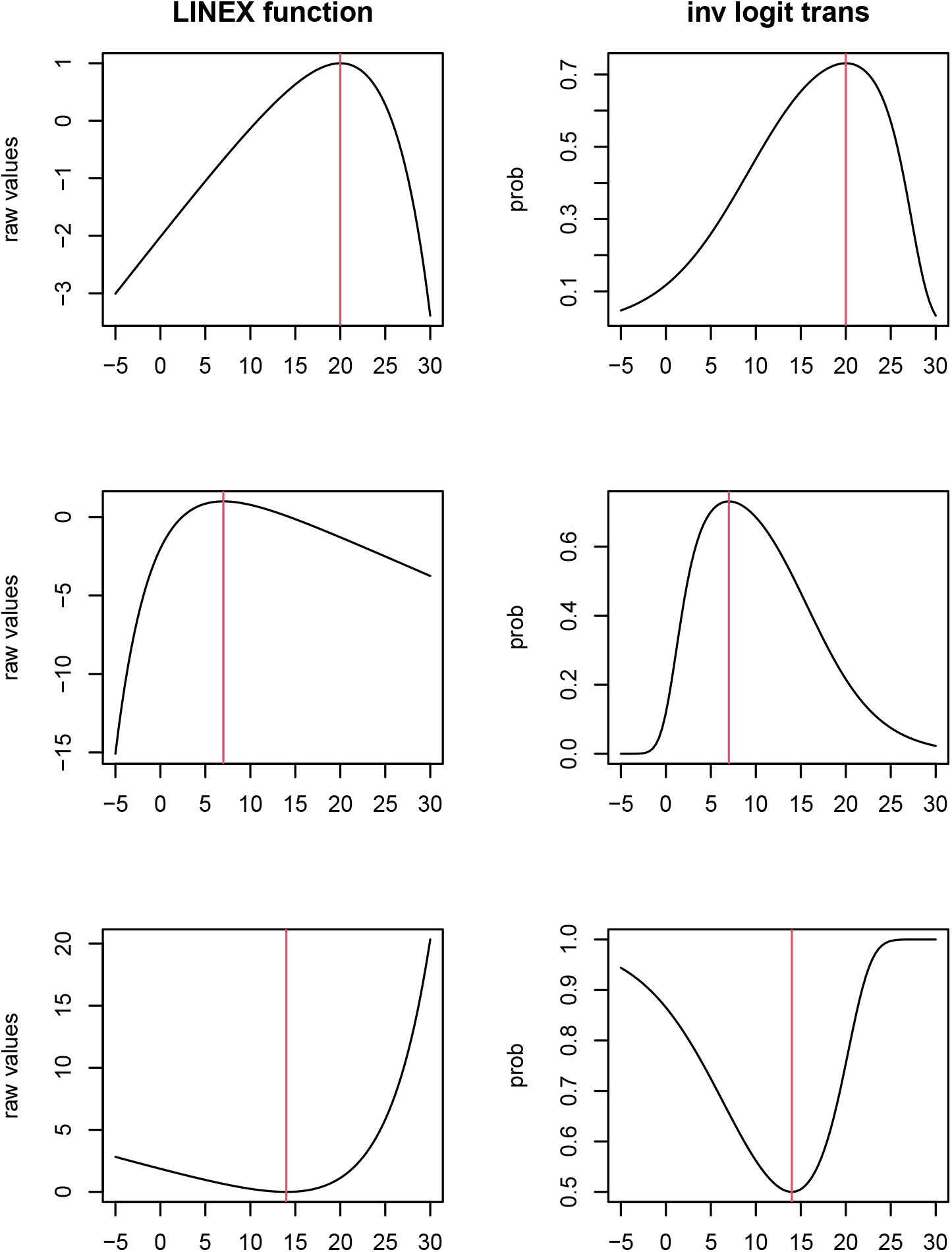
Examples of shapes that can be generated by the LINEX TPC, together with a vertical line indicating the location of the optimum. In each case, the left panel shows the shape of the non-linear predictor and the right panel shows the inverse logit transformed predictor that gives the probability of success. TOP: Left-skewed, *T*_opt_ = 20, *a* = 0.2, *c* = 2. MIDDLE: Right-skewed, *T*_opt_ = 7, *a* = −0.25, *c* = 2. BOTTOM: concave-up, *T*_opt_ = 14, *a* = 0.2, *c* = −1.

We can simulate data from this model to use for testing in a straightforward manner. We define the inverse logit transform of the LINEX function as the probability, *p*, for the target binomial distribution. We first choose parameters for the LINEX function and the number of trials in an experiment (e.g., the number of eggs). We then draw the number of “successes” (eggs that hatch) using the `rbinom` function in R with the *p* set as the value of *p* at the temperature setting, and repeat this across a set number of replicates (Fig 2). We will use these simulated data to test the inference functions in the next section.

**Figure 2.**
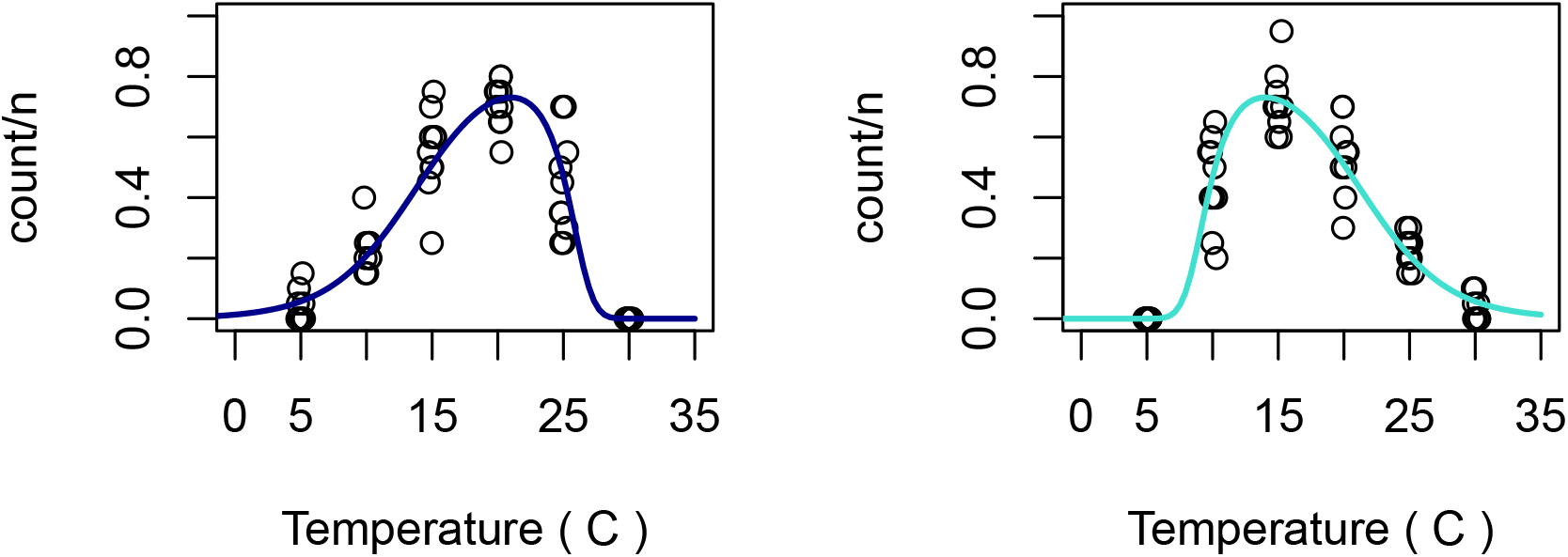
Visualization of simulated data from the LINEX TPC model, together with inverse transform of the assumed functional response (blue lines). Points correspond to 10 virtual replicates, each consisting of *n* = 20 ‘trials’ with the proportion of successes plotted (counts/*n*) at each temperature. Temperature values are jittered slightly for visualization purposes. LEFT: Left-skewed simulation, dark blue line, *T*_opt_ = 21, *a* = 0.3, *c* = 2. RIGHT: Right-skewed simulation, light blue line, *T*_opt_ = 14, *a* = −0.3, *c* = 2

### 2.3 Bayesian inference for LINEX TPC using bayesTPC

In R [12] and most other common statistical software, the sort of extensions to GLMs used here are not easy to implement by default. Typically a user would need to be able to manually specify the predictor, link, and likelihood. This must then be inputted into an appropriate optimization routine (to perform maximum likelihood estimation of parameters), or priors could additionally be specified and Bayesian estimation performed (e.g., via Markov-chain Monte Carlo). This typically requires some specialist knowledge. However, the package `bayesTPC` [13] in R allows for specification of TPCs and model formulations in a relatively straightforward way. Thus we use this package and its machinery to build a straightforward example pipeline.

Building a model in `bayesTPC` requires three steps:

1. defining a named TPC with corresponding formula (as an R “expression”)
2. defining priors for all parameters that will be fitted as a vector with named elements
3. building the model using the appropriate model specification function – here `specify_binomial_model` which assumes the appropriate logit link function and binomial likelihood.

These functions build the model in the BUGS language. This resulting model can be examined here:

~~~
{
 for (i in 1:N){
   logit(m[i]) <- (-exp(-a * (Topt - Temp[i])) + a * (Temp[i] - Topt) + cc)
   Trait[i] ∼ dbinom(m[i], n[i])
 }
cc ∼ dexp(0.1)
a ∼ dnorm(0, 1)
Topt ∼ dnorm(25, 1/1000)
}
~~~

Notice that the TPC expression is defined in terms of the temperature, which is notated as `Temp`. This is required by `bayesTPC` to ensure that all follow-on functions work properly. Parameters can otherwise be defined as the user prefers, although it is recommended that parameters not share names with functions or variables that are used elsewhere in R (for example `c` which is the concatenate function in R, or `T` and `F` which correspond to `TRUE` and `FALSE`).

## 3 Results

### 3.1 Performance on simulated data

The `bayesTPC` package requires that data be formatted as a list with elements called “Trait” (the response being modeled) and “Temp” (the corresponding temperature at which the trait is measured. Additionally, for fitting binomial models, as we are doing here, the data list should include “n”, the number of trials of which the “Trait” is the number of successes in that trial.

Once the data have been formatted and our bespoke LINEX TPC model defined, we use the built in function `b_TPC` to fit the model. There are various options that can be included. Here we specify a sampler type “AF_slice” (that works well for posteriors that may have high correlation, as TPCs often do), as well as a burn in length and number of total iterations of the MCMC to keep (for more details about possibilities see [13]). We also give an example of specifying a starting value for the sampler. Although this is not strictly necessary, it can speed up convergence.

The `bayesTPC` package includes a variety of visualization tools that utilize `coda` [14] and similar packages to allow checking of MCMC chain convergence, prior-posterior overlap, and marginal and joint posterior distributions of parameters (see Section A.1 for diagnostic plots for the model fits for the left-skewed data). It also provides summaries of model fits, and maximum *a posteriori* (MAP) estimates of parameters. For our purposes here, the most interesting output is the estimated median posterior TPC and the 95% highest posterior density (HPD) interval of this median across temperatures (Fig 3). In the simulated data set, we have a sufficient amount of data across a reasonably fine grid of temperature settings. This allows the algorithm to reconstruct the curve and parameters well.

**Figure 3.**
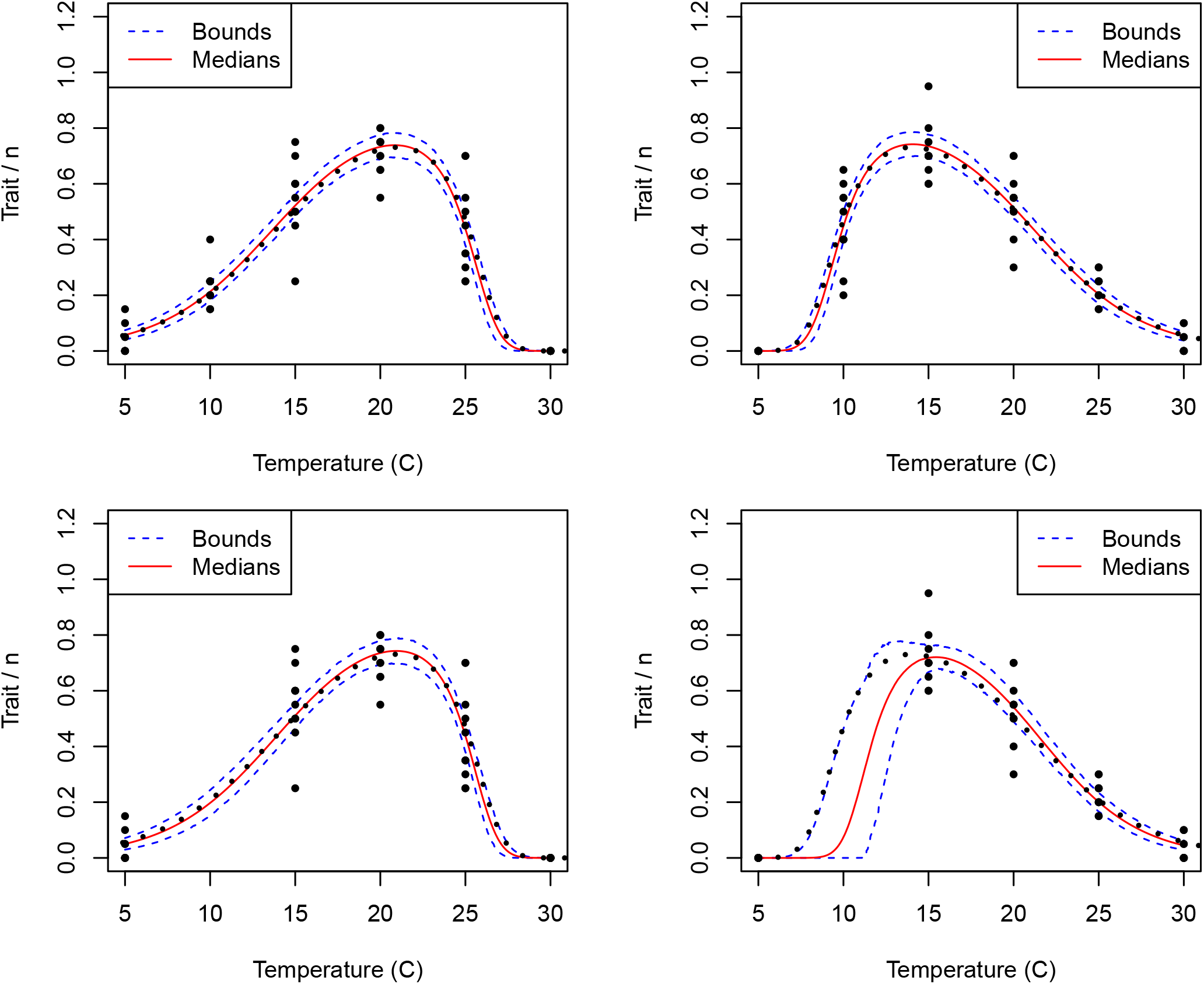
Visualization of posterior fits of the LINEX TPC to simulated data from the LINEX TPC model from the left-skewed (LEFT) and right-skewed (RIGHT) models show in Fig 2. The TOP row shows the complete data set and the BOTTOM row where the data at 10^*o*^C has been removed. The median fit is indicated with a solid red line, the 95% HPD interval of the TPC as the dashed blue lines. The true line (corresponding to the model that generated the data) is indicated with a dotted black line. Points correspond to 10 virtual replicates, each consisting of *n* = 20 ‘trials’ with the proportion of successes plotted (counts/*n*) at each temperature.

It is important to note that as the model is so flexible, it can miss the mark in cases where the experimental design (i.e., temperature settings) is less than ideal. In Fig 3 (bottom two panels) we give examples of fits obtained when data at a single temperature (here 10^°^C) are removed. For the left skewed data (Fig 3, bottom left panel), this removes data in the longer tail, which can result in two effects. First (as shown here) we can get a similar fit, with slightly increased uncertainty. This data missingness can also slow convergence and result in poor fits where the estimated skew is backwards and the peak is close to the range of the missing data (not shown). In contrast, when the data are missing closer to the peak as in the right-skewed data (Fig 3, bottom right panel) the right hand tail is still captured adequately, but the uncertainty in the location of the peak increases, which also changes the estimate of the temperature where the drop-off occurs. These are standard issues in estimation of parameters for non-linear functions if insufficient predictor settings are used, as has been noted by other authors [7].

### 3.2 Examples using ‘real’ arthropod TPC data

Next we demonstrate the performance of the LINEX TPC on real world experimental data, focusing on model selection. In some cases, theory can guide the user’s choice between fitting a symmetric or asymmetric TPC. Since the LINEX function proposed here cannot capture symmetric patterns, the user must consider assumptions and compare function choice carefully.

Here we give examples from two similar data sets where we show how the user could fit multiple functional forms and choose between them using information criteria, specifically WAIC (widely-applicable information criterion, [15]). More specifically, we explore data on the proportion of larvae/eggs that successfully develop into adults across temperatures for two species of mosquitoes, *Aedes aegypti* [16] and *Anopheles stephensi* [17].

#### 3.2.1 Larvae to adult emergence probability for *Aedes aegypti*

Our first example is from [16], one of the first studies to explore the temperature dependence of emergence probability in *Aedes aegypti*. This study observed newly emerged larvae, and recorded the proportion of these emerging as adults, at each of five fixed temperatures. Thus, the trait aggregates survival and development across multiple life stages (3 larval instars, and pupae), but does not include the probability of hatching from the egg stage. We fit the LINEX TPC in the binomial likelihood framework and a more standard quadratic GLM to these data. We use WAIC as our metric for model selection.

As we can see in Fig 4, the LINEX TPC is visually a much better fit than the quadratic – these data seem to be fairly obviously asymmetric. The WAIC scores confirm the visual assessment, showing that for these data, the LINEX TPC is strongly preferred (Fig 4).

**Figure 4.**
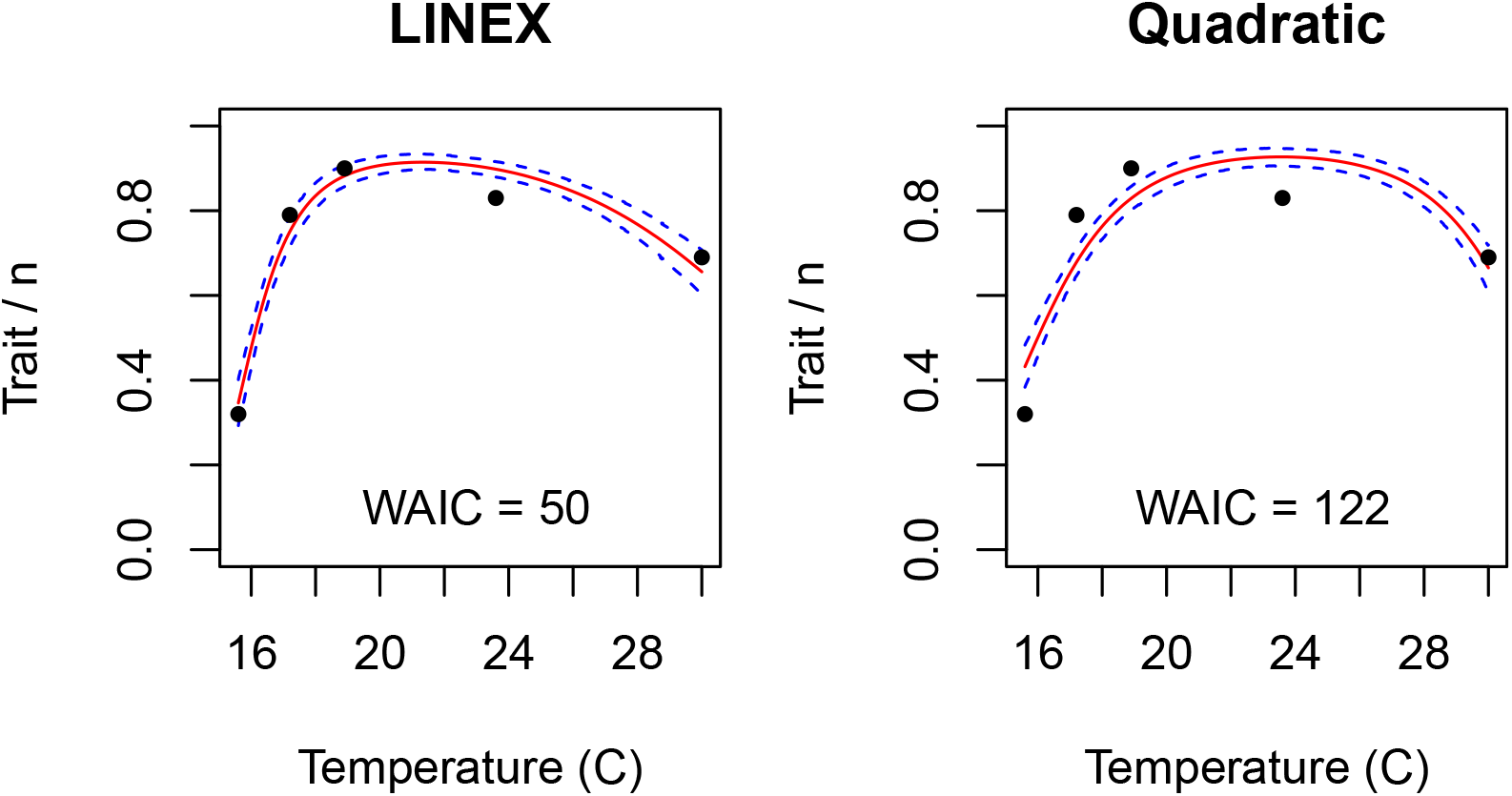
Visualization of posterior fits of the LINEX TPC (left) and quadratic generalized linear model (right) to data on probability of larvae successfully developing to the adult stage in *Aedes aegypti*across temperature (from [16]). The median fit is indicated with a solid red line, the 95% HPD interval of the TPC as the dashed blue lines. WAIC for the corresponding fits are shown in each panel

#### 3.2.2 Egg to adult emergence probability for *Anopheles stephensi*

For our second example, we examine data from [17] on the proportion of *Anopheles stephensi* eggs that successfully develop into adults across 9 temperatures and 3 different relative humidity (RH) levels. This trait aggregates survival and development across multiple life stages (egg, 3 instars, and pupa) into a single metric. TPCs for this trait are typically assumed to be symmetric (e.g., quadratic). We fit the same two models to data at each humidity level separately, and choose the best fitting model at each humidity level using WAIC.

We can see in Fig 5 that both models visually have reasonable fits at each humidity level, although the relatively flat emergence probability across a wide temperature range seen at 90% RH (bottom two rows) is not particularly well captured by either model. Model selection via WAIC prefers the symmetric quadratic function at most RH (including levels not shown here), but at 75% RH the LINEX is favored (Table 1).

**Table 1.** WAIC values for the LINEX and quadratic generalized linear model fits from Fig 5.

| RH | LINEX | QUAD |
| --- | --- | --- |
| 45 | 368.42 | 275.39 |
| 75 | 285.62 | 294.16 |
| 90 | 1007.58 | 642.67 |

**Figure 5.**
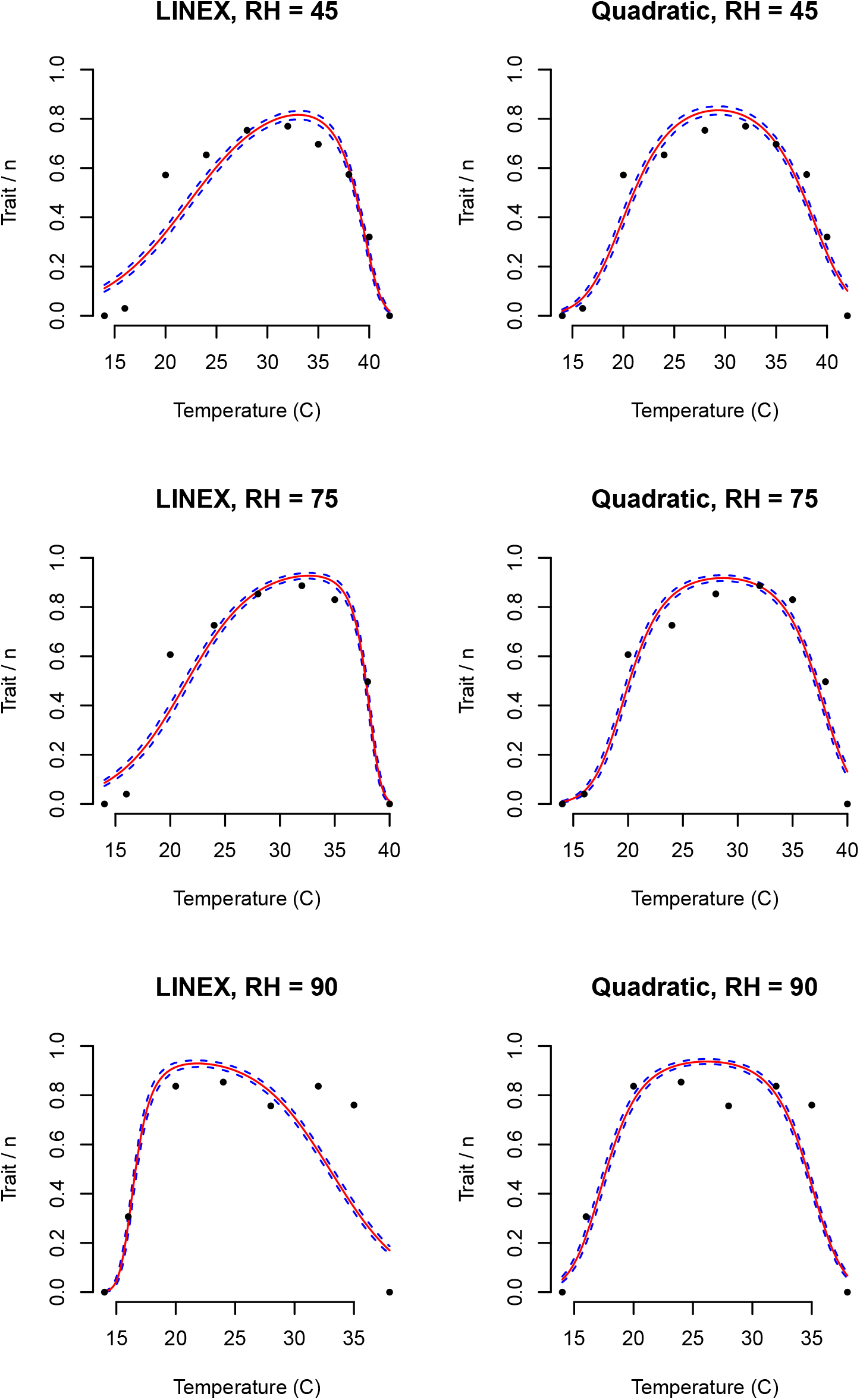
Visualization of posterior fits of the LINEX TPC and quadratic generalized linear model to data on the proportion of eggs emerging as adults across temperature for three humidity levels (from Huxley et. al, submitted). The median fit is indicated with a solid red line, the 95% HPD interval of the TPC as the dashed blue lines.

## 4 Discussion

In this paper we introduce the use of the LINEX function, a fairly simple non-linear function that can serve as an asymmetric TPC in a GLM-style modeling approach. This approach can be used model bounded data that are not well captured by a more classic piecewise continuous bounded TPC with normal (or approximately normal) noise. We showed how to implement analyses using the bayesTPC package to perform inference in a Bayesian context, exploring both the performance on simulated data (where parameters are known) and experimental data, where parameters are unknown.

Often the assumed shape of a TPC reflects assumptions about the traits. For example, compound traits, such as the probability of emergence of an adult mosquito (which integrates over multiple life stages), are frequently assumed to have symmetric TPCs. However, the choice of functions for traits are not often obvious, for example, in situations where secondary stressors may impact the trait. Thus, a practitioner usually will need to choose between functional forms when fitting models.

There are a wide variety of functions that have been proposed as TPCs in thermal ecology. These functions allow users to assume a wide variety of shapes while assuming different levels of “mechanism” that cause the shape. TPC functions also vary in their flexibility. The Briere function [18] is fairly strict, with a shape that is always left skewed (long left-hand tail), a sharp drop off, and an assumption that minimum performance must occur at temperatures above 0^°^C. In contrast, the new flexTPC [19] enables symmetric or non-symmetric shapes, left- or right-skew, and minimum temperatures of any value. The trade-off is an increase in the number of parameters needed to capture the possible shapes.

A unifying feature of these TPCs that they are explicitly bounded from below, that is the functions are ≥ 0. This boundedness is important for the vast majority of TPC applications, as the performance traits being modeled are also typically strictly ≥ 0. However, this makes established TPCs inappropriate for including within a GLM-type analysis with canonical link functions. Although it is strictly possible to include the more traditional (typically piece-wise continuous) TPCs in a GLM analysis using a unit link, this can create other problems. For example, neither the Poisson distribution nor the binomial distribution is well defined for a mean with value exactly zero. The canonical link functions explicitly force an asymptote *towards* zero to avoid this occurring. If one is instead truncating a function (e.g., a Briere) at particular values, this causes instabilities in estimation under this framework. One could instead use a small non-zero value outside of the minimum and maximum limits of the TPC. However, doing so is equivalent to assuming a minimum rate (for Poisson) or probability (for a binomial) that is constant across all other values. This is both a biologically poor assumption, and one that can bias estimation of the TPC and predictions from the model.

The LINEX TPC, together with a simple quadratic (to capture symmetric responses), are able to represent a wide range of shapes that are appropriate to use within the logistic model framework because they have infinite support – that is they can produce responses from −∞ to ∞. Further, because of this property, they are also appropriate to model other sorts of data within the GLM-flavor framework. For example, count data (such as may be recorded for fecundity) could be modeled with the LINEX TPC and the log-link to either a Poisson or negative-binomial likelihood. Thus, this relatively simple three-parameter function can be used for a wide variety of thermal trait (or other) data.

Although the model is fairly simple, due to the non-linearity in the parameters, the choice of priors for Bayesian inference can still be a bit of an art. The temptation is often to choose priors that are wide and relatively un-informative for all parameters, but this can result in inadequate fitting and bias. For example, if one lets both *c* and *a* have wide priors, particularly if both can be either positive or negative, the resulting fit can be biased. The rate of convergence can be greatly improved by instead choosing weakly informative priors for one or both of these parameters. For instance, if we know that our peak probability is high-ish – above 0.3 or so – then a prior that constrains *c* to be positive improves the inference substantially. Further, if one has biological knowledge about the likely direction of the skew of the trait, constraining *a* to reflect this will also improve inference. In the examples given in this study, we chose weakly informative priors to substantially improve convergence. Practitioners taking a maximum likelihood approach would likely need to constrain optimization routines in similar ways to obtain good results.

## Appendix

### A.1 Diagnostics and summary for fit of left-skewed simulation example

**Figure A1.**
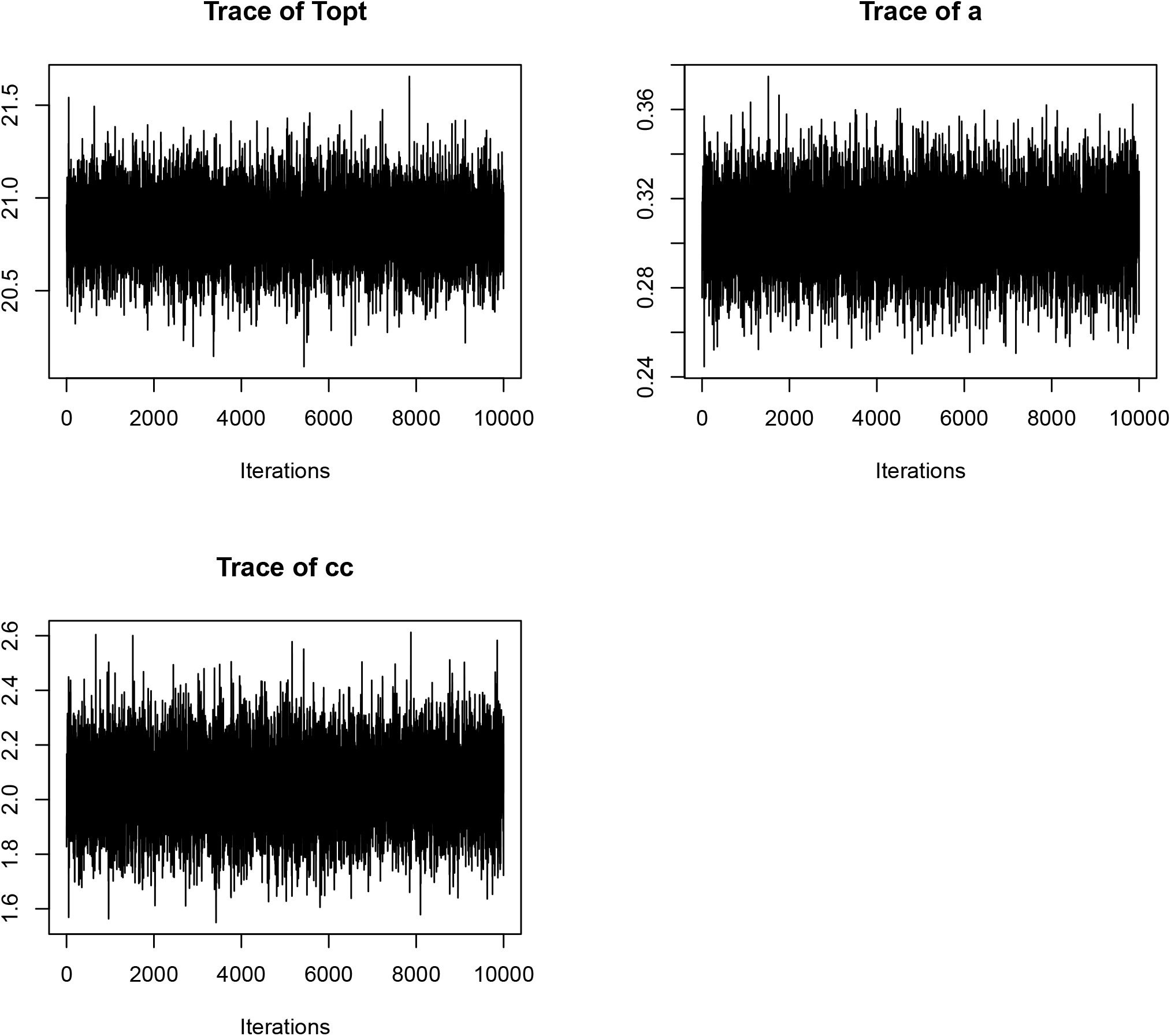
Trace plots for the LINEXP model fit to the left skewed simulation data.

**Figure A2.**
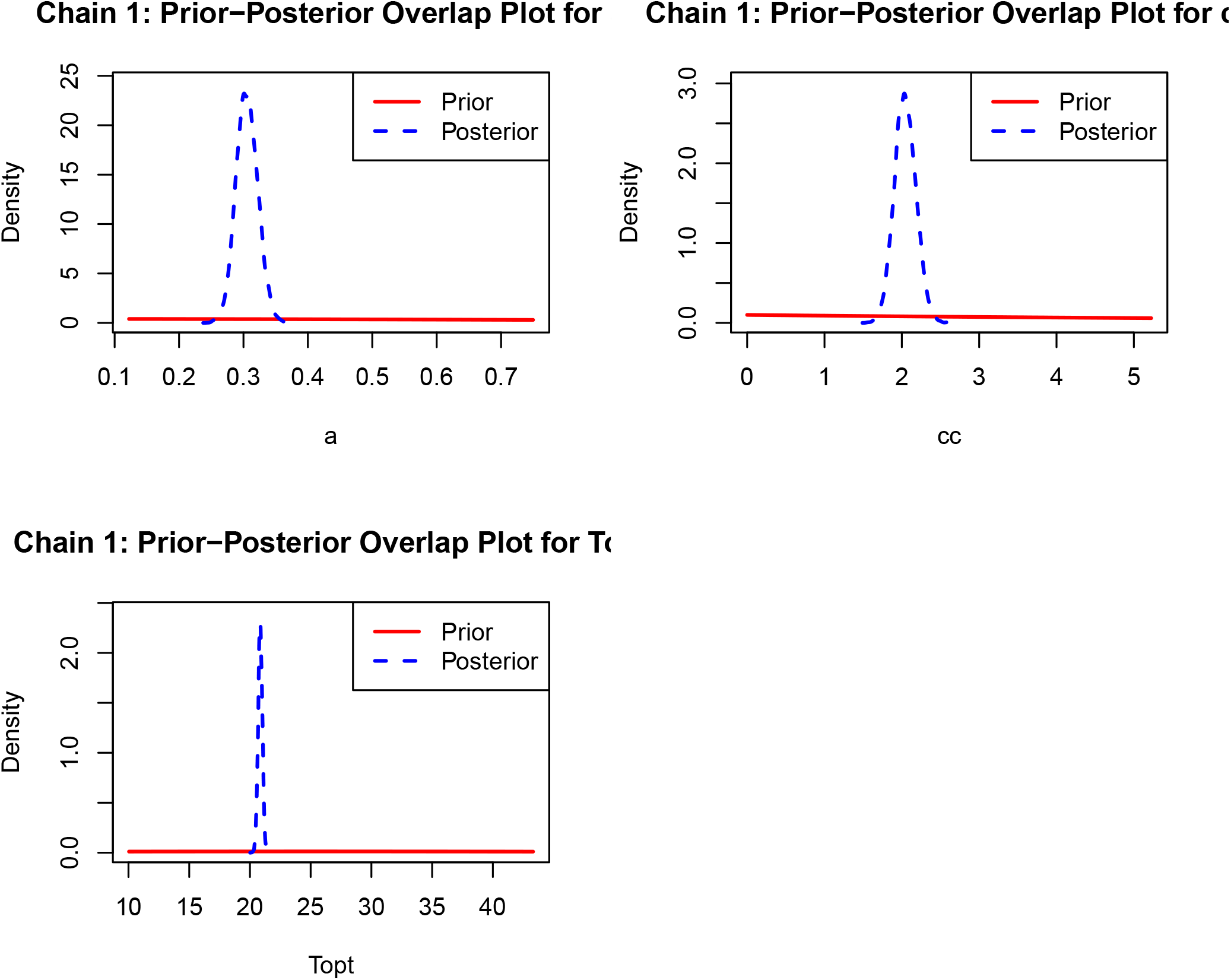
Prior-posterior overlap plots for the LINEXP model fit to the left skewed simulation data.

**Figure A3.**
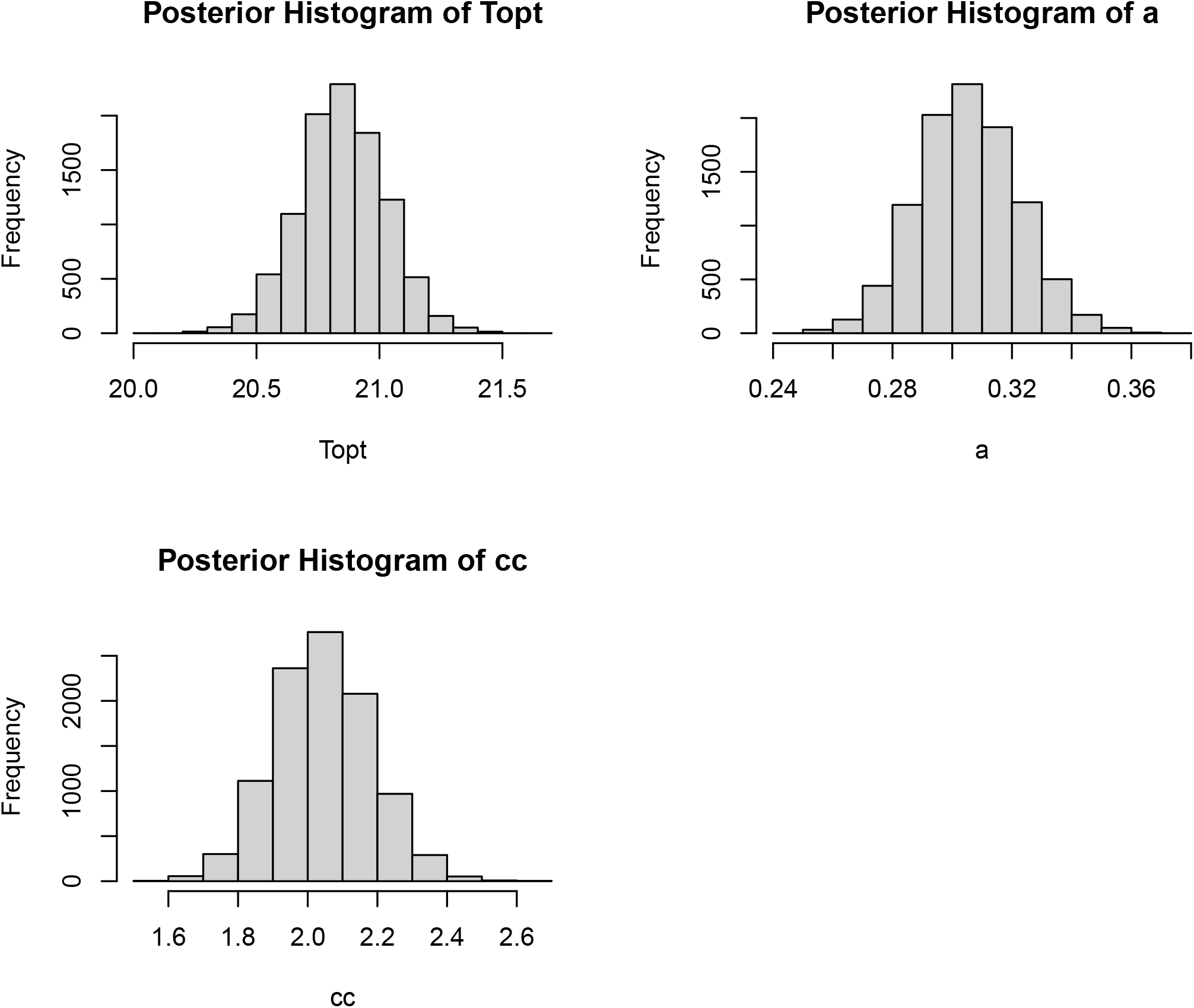
Histograms of the marginal posterior samples of parameters for the LINEXP model fit to the left skewed simulation data.

**Figure A4.**
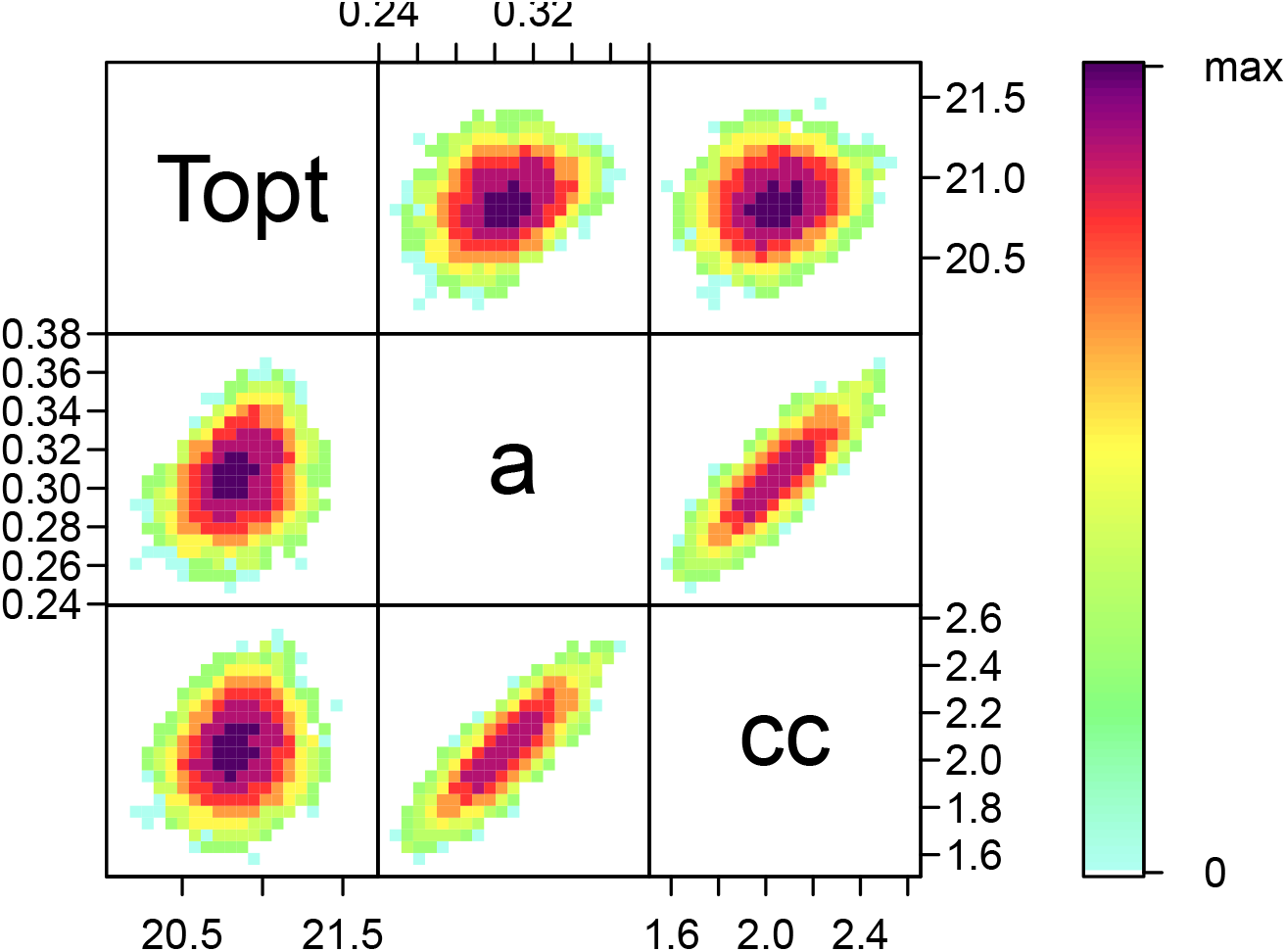
Pairs plot of samples from the joint posterior of parameters for the LINEXP model fit to the left skewed simulation data.

~~~
bayesTPC MCMC of Type:
 LinExp_binom
Formula:
 logit(m[i]) <- (-exp(-a ^*^ (Topt - Temp)) + a ^*^ (Temp - Topt) + cc)
Distribution:
 Trait[i] ∼ dbinom(m[i], n[i])
Priors:
 cc ∼ dexp(0.1)
 a ∼ dnorm(0, 1)
 Topt ∼ dnorm(25, 1/1000)
Max. A Post. Parameters:
   Topt a cc log_prob
 20.8308 0.3059 2.0484 -110.2263
MCMC Results:
Iterations = 1:10000
Thinning interval = 1
Number of chains = 1
Sample size per chain = 10000
1. Empirical mean and standard deviation for each variable,
  plus standard error of the mean:
         Mean     SD       Naive SE Time-series SE
Topt 20.8495  0.17646    0.0017646   0.0018403
a     0.3054  0.01684    0.0001684   0.0001822
cc    2.0439  0.13906    0.0013906   0.0014876
2. Quantiles for each variable:
           2.5%       25%       50%       75%     97.5%
Topt    20.5008   20.7352   20.8486   20.9687   21.1932
a        0.2733    0.2939    0.3052    0.3168    0.3392
cc       1.7783   1.9501    2.0400    2.1365    2.3190
~~~

## Notes

### Competing Interest Statement

The authors have declared no competing interest.

### Funding

- U.S. National Science Foundation: DBI #2016264

## References

1. Angilletta Jr MJ, Niewiarowski PH, Navas CA. The evolution of thermal physiology in ectotherms. Journal of thermal Biology. 2002;27: 249–268.

2. Huey RB, Stevenson R. Integrating thermal physiology and ecology of ectotherms: A discussion of approaches. American Zoologist. 1979;19: 357–366.

3. Schoolfield RM, Sharpe P, Magnuson CE. Non-linear regression of biological temperaturedependent rate models based on absolute reaction-rate theory. Journal of theoretical biology. 1981;88: 719–731.

4. Amarasekare P, Savage V. A framework for elucidating the temperature dependence of fitness. The American Naturalist. 2012;179: 178–191.

5. Gajewski Z, Stevenson LA, Pike DA, Roznik EA, Alford RA, Johnson LR. Predicting the growth of the amphibian chytrid fungus in varying temperature environments. Ecology and Evolution. 2021;11: 17920–17931.

6. Padfield D, O’Sullivan H, Pawar S. rTPC and nls. Multstart: A new pipeline to fit thermal performance curves in r. Methods in Ecology and Evolution. 2021;12: 1138–1143.

7. Pawar S, Dell AI, Savage VM, Knies JL. Real versus artificial variation in the thermal sensitivity of biological traits. The American Naturalist. 2016;187: E41–E52.

8. Parsian A, Kirmani S. Estimation under LINEX loss function. Handbook of applied econometrics and statistical inference. CRC Press; 2002. pp. 75–98.

9. Singh U, Gupta PK, Upadhyay S. Estimation of exponentiated weibull shape parameters under LINEX loss function. Communications in Statistics-Simulation and Computation. 2002;31: 523–537.

10. Li X, Shi Y, Wei J, Chai J. Empirical bayes estimators of reliability performances using LINEX loss under progressively type-II censored samples. Mathematics and Computers in Simulation. 2007;73: 320–326.

11. Doostparast M, Ahmadi MV, Ahmadi J. Bayes estimation based on joint progressive type II censored data under LINEX loss function. Communications in Statistics-Simulation and Computation. 2013;42: 1865–1886.

12. R Core Team. R: A language and environment for statistical computing. Vienna, Austria: R Foundation for Statistical Computing; 2025. Available: https://www.R-project.org/

13. Sorek S, Smith Jr. JW, Huxley PJ, Johnson LR. ‘bayesTPC’: Bayesian inference for thermal performance curves in r. Methods in Ecology and Evolution. 2025;16: 687–697. doi:10.1111/2041-210X.70004

14. Plummer M, Best N, Cowles K, Vines K. CODA: Convergence diagnosis and output analysis for MCMC. R News. 2006;6: 7–11. Available: https://journal.r-project.org/archive/

15. Gelman A, Hwang J, Vehtari A. Understanding predictive information criteria for bayesian models. Statistics and Computing. 2013;24.

16. Headlee TJ. A continuation of the studies of the relative effects on insect metabolism of temperature derived from constant and varied sources. Journal of Economic Entomology. 1942;35: 785–786.

17. Huxley PJ, Brown JJ, St. Laurent B, Johnson B, Cheung OY, Asamoah A, et al. Beyond temperature: Relative humidity systematically shifts the temperature dependence of population growth in a malaria vector. bioRxiv. 2025; 2025–05.

18. Briere J-F, Pracros P, Le Roux A-Y, Pierre J-S. A novel rate model of temperature-dependent development for arthropods. Environmental Entomology. 1999;28: 22–29.

19. Cruz-Loya M, Mordecai EA, Savage VM. A flexible model for thermal performance curves. bioRxiv. 2024.

